# Dropout in Neural Networks Simulates the Paradoxical Effects of Deep Brain Stimulation on Memory

**DOI:** 10.1101/2020.05.01.073486

**Authors:** Shawn Zheng Kai Tan, Richard Du, Jose Angelo Udal Perucho, Shauhrat S. Chopra, Varut Vardhanabhuti, Lee Wei Lim

## Abstract

Neuromodulation techniques such as Deep Brain Stimulation (DBS) are a promising treatment for memory-related disorders including anxiety, addiction, and dementia. However, the outcome of these treatments appears to be paradoxical, as the use of these techniques can both disrupt and enhance memory even when applied to the same brain target. In this paper, we hypothesize that disruption and enhancement of memory through neuromodulation can be explained by the dropout of engram nodes. We used a convolutional neural network to classify handwritten digits and letters, applying dropout at different stages to simulate DBS effects on engrams. We showed that dropout applied during training improves the accuracy of prediction, whereas dropout applied during testing dramatically decreases accuracy of prediction, which mimics enhancement and disruption of memory, respectively. We further showed that transfer learning of neural networks with dropout had increased accuracy and rate of learning. Dropout during training provided a more robust “skeleton” network where transfer learning can be applied, mimicking the effects of chronic DBS on memory. Overall, we show that dropout of nodes can be a potential mechanism by which neuromodulation techniques such as DBS can both disrupt and enhance memory and provides a unique perspective on this paradox.

## Introduction

Memory systems are crucial for survival and, to a large extent, define who we are. However, memory systems can fall into disease when expressed pervasively (e.g., anxiety or addiction) or degenerated (e.g., dementia)–both of which are major health challenges worldwide [1. 2]. Neuromodulation techniques such as Deep Brain Stimulation (DBS) have shown promising results as treatments for such memory-related disorders [3. 4], yet the mechanisms behind these effects are still largely unknown. Furthermore, the effects of treatments such as DBS appear to be paradoxical, in that they can both disrupt [5. 6] and enhance memories [7. 8], even when applied to the same brain target [3]. We have previously suggested that DBS is able to disrupt memory by “removing” nodes in an engram [4]. However, this theory remains untested partly due to the lack of technology to monitor large engram networks in real-time. Besides, this theory does not explain (at least directly) how memory enhancement is achieved.

The development of machine learning techniques offers a unique computational approach to simulate hypothetical models of learning and memory, and the effects of manipulation on memory, which we have previously used to highlight potential mechanisms of memory disruption by DBS [4]. To model the learning process, we trained a convolutional neural network (CNN) to classify handwritten digits and letters. CNNs are a type of artificial neural network that are commonly applied to image recognition tasks. The concept of CNNs was developed from early observations of the visual cortex, in which groups of neurons fired distinctively in response to different light patterns (e.g., straight lines, circles). In a typical CNN, the network extracts features using a convolutional layer followed by classification of the features. For image tasks, this convolutional layer is comprised of a series of convolution filters that are associated with particular patterns of pixels, which mimics receptive fields in the retina. These trainable filters are also referred to as weights of the network, similar to synapse/synaptic strength in biological systems. In image classification, the CNN decomposes the input image into patterns of pixels known as features. First, an input image is partitioned into non-overlapping regions, with each region mapped to a specific neuron. Second, the neurons are convolved by multiple filters to generate a feature map in the convolution layer. The resultant feature maps can be further decomposed by inputting these maps into successive convolution layers. After a specified number of decompositions, the resultant features are used to classify the input image using the fully connected layer.

In this paper, we hypothesize that the paradoxical ability of DBS to both disrupt and enhance memory can be explained through dropout (a process of randomly shutting down or dropping neurons) in engram nodes by using CNN to simulate learning and memory.

## Methods

### Dataset

To model the learning process, we trained a convolutional neural network (CNN) to classify handwritten digits and letters in the EMNIST dataset. The EMNIST dataset is a public database of over 800,000 handwritten digits and letters across 62 different classes [9]. In our study, we used the EMNIST balanced dataset, which is derived by merging similar classes of letters. This dataset contains 131,600 28×28 pixel images of 47 balanced classes (10 digits, and 37 uppercase and lowercase letters).

### Network Architecture

The CNN consist of an input layer of size 28×28×1 followed by two convolution layers of 32 filters and 64 filters, leading to a feature map of size 5×5×64. No padding was used for the convolutions, and a filter of size 3×3 was used in both layers. At the end of each convolution layer, we applied max pooling with a 2×2 window. Max pooling is a standard process for reducing the dimensions of feature maps by sampling the maximum value for a given window size, which forces the network to enhance and focus on important features [10]. Global max pooling was applied to the feature map to extract 64×1 latent features. For feature classification, the features were passed through to a fully connected layer of 100 neurons. A rectified linear unit (ReLU) was applied as an activation function in all the layers, which is a ramp function where all negative value neurons are zeroed to ensure unidirectionality. The parameters in all the networks were optimised using binary cross-entropy or loss function. All networks were trained for 500 epochs in batches of 10 images. An Epoch is defined as one complete run-through of all the training data. For the analysis, the average loss of training data and average classification accuracy of the testing data were evaluated at the end of each epoch.

### Experiments

We conducted two sets of experiments in this study. In experiment 1, we trained a network to classify 10 different digits (0 to 9) from the EMNIST balanced dataset. We applied dropout (a process of randomly shutting down or dropping neurons) either during the training stages, or just prior to testing at each epoch. We used a dropout rate of 50%, such that half the neurons were dropped at each step (Fig 1A,B). Dropout was only done on fully connected latter layers of the network. As the aim of the study was not to train a network to classify the digits accurately but to analyse the learning process, we randomly sampled 1000 digits as the training dataset and the network was evaluated on another 1000 randomly sampled digits as the testing dataset. All networks were trained with the same training dataset and tested with the same testing dataset.

In experiment 2, we transferred the network to learn 26 uppercase letters (A to Z) in a process known as transfer learning, in which the networks and weights of the control (non-dropout) group and dropout group were applied to the new task (Fig 1C). We retrained the network to recognise the uppercase letters by stripping the last output layer and replacing it with the 26 classes corresponding to each letter class. We evaluated the performance of the transfer learning using the trained network with dropout and without dropout, and the performance of a network trained with letters from scratch without transfer learning (Fig 1C). Due to the increased complexity of more classes, we used 5000 randomly sampled letters as the training dataset and 1000 randomly sampled letters as the testing dataset.

**Figure 1.**
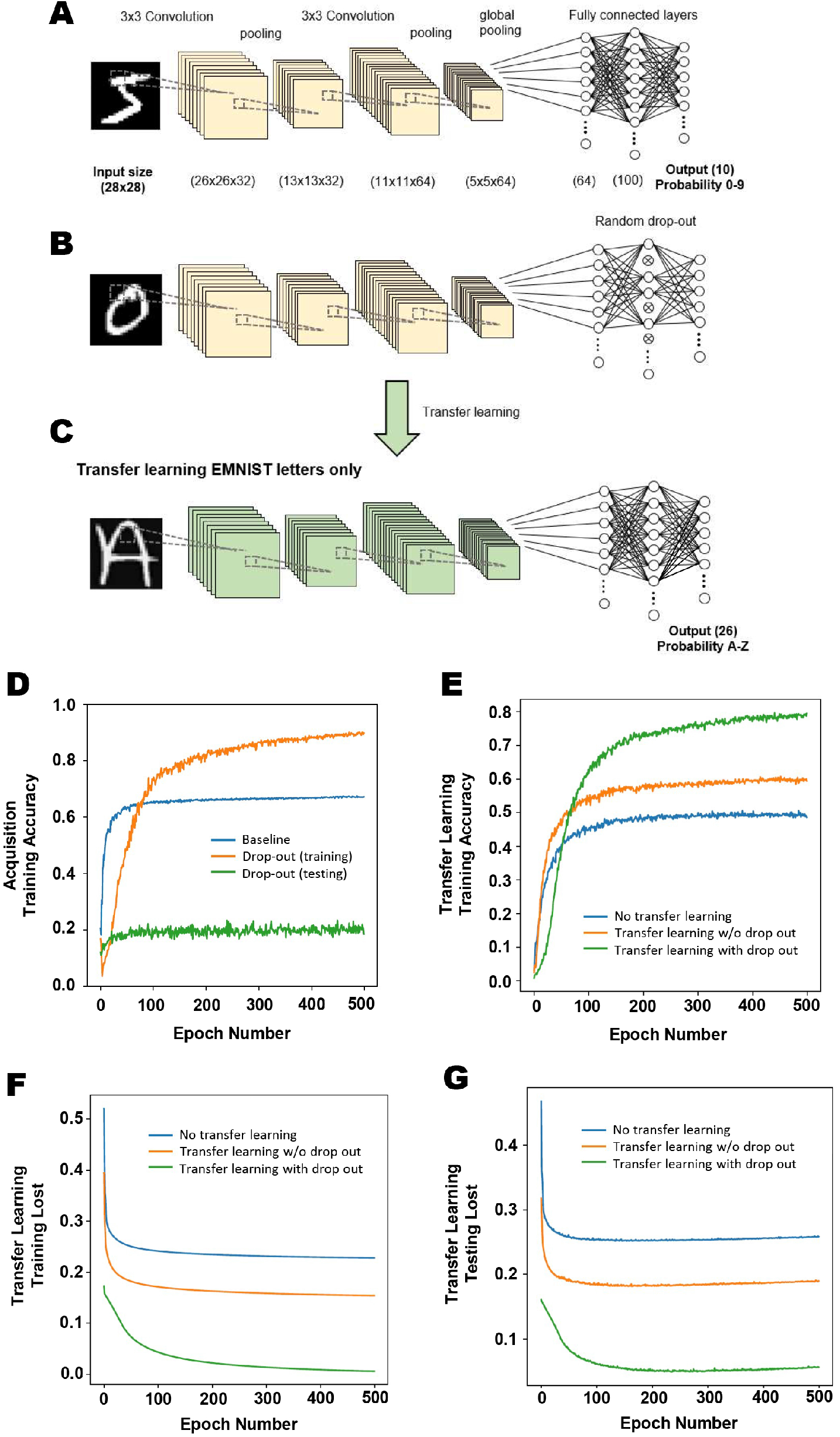
Dropout in neural nodes disrupt or enhance learning in neural networks depending on when it is applied. A convolutional neural network (CNN) was trained to classify handwritten digits in the EMNIST dataset (A). Dropout was applied to 50% of nodes in the fully connected layer (B). Transfer learning was then done, and the network retrained in recognising uppercase letters (C). Dropout applied during training improves prediction accuracy, whereas dropout applied during testing dramatically decreases prediction accuracy (D). Compared to non-dropout networks, transfer learning of neural networks with dropout had increased accuracy (Fig 1E) and had lowered training loss (F) and testing loss (G).

## Results

### Experiment 1

To simulate the effects of DBS on memory, we applied dropout at either the training or testing stages. Accuracy of prediction was used as an indication of how “well” the neural network had learnt the task, which serve as a proxy for memory. Therefore, a higher accuracy of prediction should indicate higher memory function. In our experiment, dropout applied during training improves the accuracy of prediction, whereas dropout applied during testing dramatically decreases the accuracy of prediction (Fig 1D).

### Experiment 2

To simulate a novel memory task post chronic DBS, we applied a process of transfer learning on the trained network, in which the networks and weights of the control (non-dropout) group and dropout groups were then applied to a new task. Overall, transfer learning of the network showed increased accuracy compared to a network without transfer learning (Fig 1E), indicating higher memory function. In addition, there were decreased testing and training loss (Fig 1F,G), with the loss defined as the average errors made across the testing or training data, respectively. Decreased loss indicates better performance or better fitting to the training/testing data, which can therefore serve as a proxy for increased rate of learning. Transfer learning in neural networks with dropout resulted in better accuracy (Fig 1E) and lower training and testing loss (Fig 1F,G) compared to transfer learning in networks with no dropout.

## Discussion

We have previously argued that timing plays an important role in the outcome of DBS on memory [3]; DBS applied post or during behaviour testing tended to disrupt memory, whereas DBS applied prior to behaviour testing tended to enhance memory. However, the mechanisms behind these outcomes are still relatively unknown. Indeed, we found that dropout applied during training improves the accuracy of prediction, which is similar to the enhancement of memory seen when DBS was applied prior to behaviour testing in our previous animal studies [8]. On the contrary, we found that dropout applied post training dramatically decreases the accuracy of prediction, which is similar to the disruption of memory seen when DBS was applied during consolidation of memory in our previous animal studies [6](Fig 1D).

We used CNNs (as opposed to a standard fully connected networks) in this study as they are commonly applied in image recognition tasks. Although CNNs are modelled on the visual cortex, dropout was only applied on fully connected latter layers in the network to represents dropouts applied in the hippocampus rather than the visual cortex, which mimic our previous experiments on the effects of prefrontal cortex stimulation on the hippocampus [6. 8].

One divergence from animal experiments emerges from experiment 1 on the enhancement of memory as compared to our previous experiments - chronic DBS was not applied during the training of a task, but rather in home-cage prior to behaviour experiments. To more accurately represent the enhancement of memory through chronic DBS prior to a memory task [8], we applied transfer learning in previously trained networks with or without dropout. We showed that transfer learning did indeed increase the accuracy of prediction and decreased training and testing loss (representing an increased learning rate), indicating that transfer learning was successful in this model. More importantly, we showed that transfer learning applied to neural networks with dropout increased the accuracy of prediction (representing higher memory function) as compared to neural networks without dropout. We further showed that transfer learning applied to neural networks with dropout had lower training and testing loss, indicating not only improved memory function, but also increased rate of learning. Overall, we showed that applying dropout during training provides a more robust “skeleton” network and applying transfer learning in this network increased accuracy and decreased training and testing loss. This model showed similar results to our previous study of chronic mPFC DBS on memory enhancement [8] and suggests a potential mechanism for this process. An early hypothesis of the mechanism of DBS was that it creates a temporary neural activity lesion, and we further showed that prelimbic cortex DBS was associated with the disruption of memory and a drop in the neural activity marker c-fos in the ventral hippocampus [6]. Dropout of neural nodes, while congruent with the disruption of memory, has not played a major role in the mechanistic understanding of memory enhancement seen in DBS (though it should be noted that other mechanism such as neurogenesis and wave syncing have been suggested [3]). In this paper, we propose that DBS causes dropout in neural nodes that “forces” the activation of new pathways and creates more robust networks, similar to how dropout enhanced the neural networks. To the best of our knowledge, this paper is the first to suggest a cohesive mechanism in which disruption of neural activity through DBS can lead to both disruption and enhancement of memory. Although no behavioural experiments were performed in conjunction with our modelling, we based the models on our previous animal behavioural experiments that showed memory enhancement with DBS prior to the behavioural task [8] and memory disruption with DBS during consolidation of memory [6], which demonstrated similar results to the present paper.

In conclusion, using a machine learning model based on previous animal experiments, we showed that dropout of nodes can be a potential mechanism in which neuromodulation techniques like DBS can both disrupt and enhance memory. This paper serves as a basis for further experimentation on engrams in understanding the effects of neuromodulation on memory

## Acknowledgements & Disclosures

The scientific work was funded by grants from the Hong Kong Research Grant Council (RGC-ECS 27104616) and The University of Hong Kong URC Supplementary Funding (102009728) awarded to LWL. All authors declare no competing financial interests or potential conflicts of interest.

## Notes

### Competing Interest Statement

The authors have declared no competing interest.

